# ppGpp regulates NAD-RNA decapping by *Escherichia coli* NudC and RppH

**DOI:** 10.64898/2026.11.19.767904

**Authors:** Yogeshwari Singh, Lingting Li, Raymond George, Yu Zhang, Jeremy G. Bird

**Affiliations:** Department of Biological Sciences, University of Delaware; Key Laboratory of Synthetic Biology, Institute of Plant Physiology and Ecology, Chinese Academy of Sciences, China

**Keywords:** RNA turnover, RNA degradation, crystallography, RNA metabolism, RNA-protein interaction, NAD-capped RNA, RNA decapping, ppGpp, Nudix hydrolase, RNA stability

## Abstract

The regulatory landscape of RNA includes diverse non-canonical 5’ cap structures, including Nicotinamide Adenine Dinucleotide (NAD^+^). In *E. coli*, the NAD cap and its cognate decapping enzyme NudC have provided important insight into NAD^+^-capped RNA stability; however, broader regulatory control of NAD-capped RNA degradation has remained unclear. Here, we demonstrate that the *E. coli* Nudix hydrolase RppH, known to act on 5’ triphosphorylated RNA, also functions as an NAD^+^-decapping enzyme. RppH removes NAD^+^ caps from both linear and structured (hairpin) RNA substrates, revealing a previously unrecognized role in NAD^+^-RNA metabolism. These findings suggest that RppH may function alongside NudC to regulate the stability of NAD-capped transcripts. We further establish a direct relationship between NAD-RNA turnover and the alarmone ppGpp. Quantitative inhibition assays demonstrate that ppGpp inhibits decapping of NAD^+^-RNA by both NudC and RppH and indicate coordinated regulation of NAD-RNA decay during stress conditions. To provide a structural basis for these observations, we determined crystal structures of NudC bound to NADH and ppGpp. These structures reveal that ppGpp occupies the NudC active site, supporting a competitive inhibition mechanism. Together, our findings identify RppH as a novel NAD^+^-decapping enzyme and establish ppGpp as a regulator of NAD-RNA decay pathways, linking RNA capping and metabolic stress signaling.

## Introduction

The 5’ end of RNA plays a critical role in RNA stability, processing, and translation. Eukaryotic RNA has long been known to be modified at the 5’-end by the canonical 7-methylguanosine (mTG) cap. However, the discovery of non-canonical RNA caps derived from primary metabolites has expanded the RNA cap landscape (1–3). The most well-studied non-canonical cap is nicotinamide adenine dinucleotide (NAD), first identified in bacteria and subsequently observed across all domains of life, including in yeast, plants, humans, archaea, mitochondria, bacteriophages, and other viruses(1–10). In *Escherichia coli*, oxidized NAD (NAD^+^) caps have been identified on a subset of small regulatory RNAs and mRNAs (4). This 5’-NADylation enhances the stability of these transcripts and notably, the proportion of NAD^+^-capped RNAs increases under conditions associated with a high NAD^+^-to-ATP ratio, such as cellular stress and stationary phase(4, 11, 12). These findings together suggest that changes in NAD^+^ capping may provide a dynamic mechanism for modulating RNA stability.

*E. coli* has multiple enzymes that can process RNA 5’-ends. Cleavage of the RNA 5’-end to expose a monophosphate is the first step in RNase E-mediated RNA decay (13–15). RNase E preferentially interacts with the 5’-monophosphorylated end of substrate RNAs and internally cleaves the RNA into short fragments that are handled by other nucleases(13–16). *E. coli* RppH is a well-studied RNA pyrophosphohydrolase that has been demonstrated to primarily convert 5’-triphosphorylated RNA to 5’-monophosphorylated RNA with some activity on other substrates including dinucleotide polyphosphates and UDP-sugars (15, 17). NudC is an NADH hydrolase that is the primary NAD cap decapping enzyme, as it strongly prefers to act on RNA 5’-NAD modifications over free NAD in the cell, in addition to removing Coenzyme A as well (18–20). When NudC hydrolyzes NAD, it releases nicotinamide mononucleotide (NMN) and generates a monophosphorylated 5’-end (18, 21). The 5’-monophosphorylated RNA products of both RppH and NudC are the primary substrates of RNase E, a 5’-end dependent endonuclease whose activity constitutes the first step of the RNA decay pathway (15, 22). Understanding the regulation of cap removal by decapping enzymes is essential for defining the function of RNA caps in regulating degradation.

Here, we expand the regulatory landscape surrounding RNA cap stability by showing that *E. coli* RppH can remove NAD^+^ caps from RNA, consistent with the activity of its homolog from *Bacillus subtilis* (23). Moreover, we find that the bacterial alarmone ppGpp inhibits the decapping activity of both NudC and RppH at physiological concentrations. NudC has recently been identified in a screen to identify interactors of ppGpp and ppGpp has been demonstrated to inhibit RppH activity on nucleotide 5’-triphosphate ends (17, 24). ppGpp-mediated regulation of RNA degradation has previously been proposed in *Streptomyces* species, where ppGpp inhibits the activity of Polynucleotide Phosphorylase (PNPase), but does not interact with *E. coli* PNPase (25). Our crystal structures of NudC bound to NADH and ppGpp reveal that ppGpp directly competes with cap binding. By implicating ppGpp as a direct regulator of NAD-RNA turnover, our work provides a potential link between bacterial stress responses and cap-dependent RNA stability. Based on the mechanism by which these enzymes interact with ppGpp, we propose that the alarmone may regulate Nudix family enzymes broadly by binding to their active centers acting as competitive inhibitors.

## Results

### RppH contributes to 5’ NAD^+^ cap cleavage and RNA degradation

RppH is a member of the Nudix hydrolase enzyme superfamily and has been primarily characterized as an RNA pyrophosphohydrolase responsible for removing 5’-triphosphates from RNA (14, 15). Given its catalytic mechanism, and the ability of homologs from other bacteria to act on NAD^+^-capped RNAs, we tested whether *E. coli* RppH can also process NAD-capped RNA substrates. We evaluated decapping ability by incubating RppH with ^32^P-radiolabeled hairpin or linear NAD^+^-RNA. Examination of the resulting RNA species by urea-PAGE showed RppH-dependent accumulation of a decapped RNA product, demonstrating that RppH can remove the NAD^+^ cap from hairpin and linear NAD^+^-RNA substrates efficiently (Fig. 1). RppH displayed greater activity toward linear RNA, reaching ∼80% decapping at the highest enzyme concentration, compared with ∼60% for the structured hairpin substrate. These results suggest that RNA secondary structure reduces substrate NAD accessibility for RppH, consistent with its previously reported preference when acting on 5’-triphosphate RNA substrates (26). Whereas previous work reported that *E. coli* RppH did not decap NAD(H)-capped dinucleotide RNA (11), our results demonstrate that RppH decaps longer NAD^+^-capped RNA substrates, including both 32-nucleotide hairpin and 40-nucleotide linear RNA (Fig. 1). It is likely that longer RNA substrates bind to the established RNA-binding pocket adjacent to the enzyme’s active site and correctly position the NAD in the substrate binding pocket to allow cleavage of the diphosphate linkage. While RppH was less active than NudC under similar assay conditions, our results nevertheless establish that RppH is a second *E. coli* enzyme with the ability to remove NAD^+^ caps from RNA.

**Figure 1.**
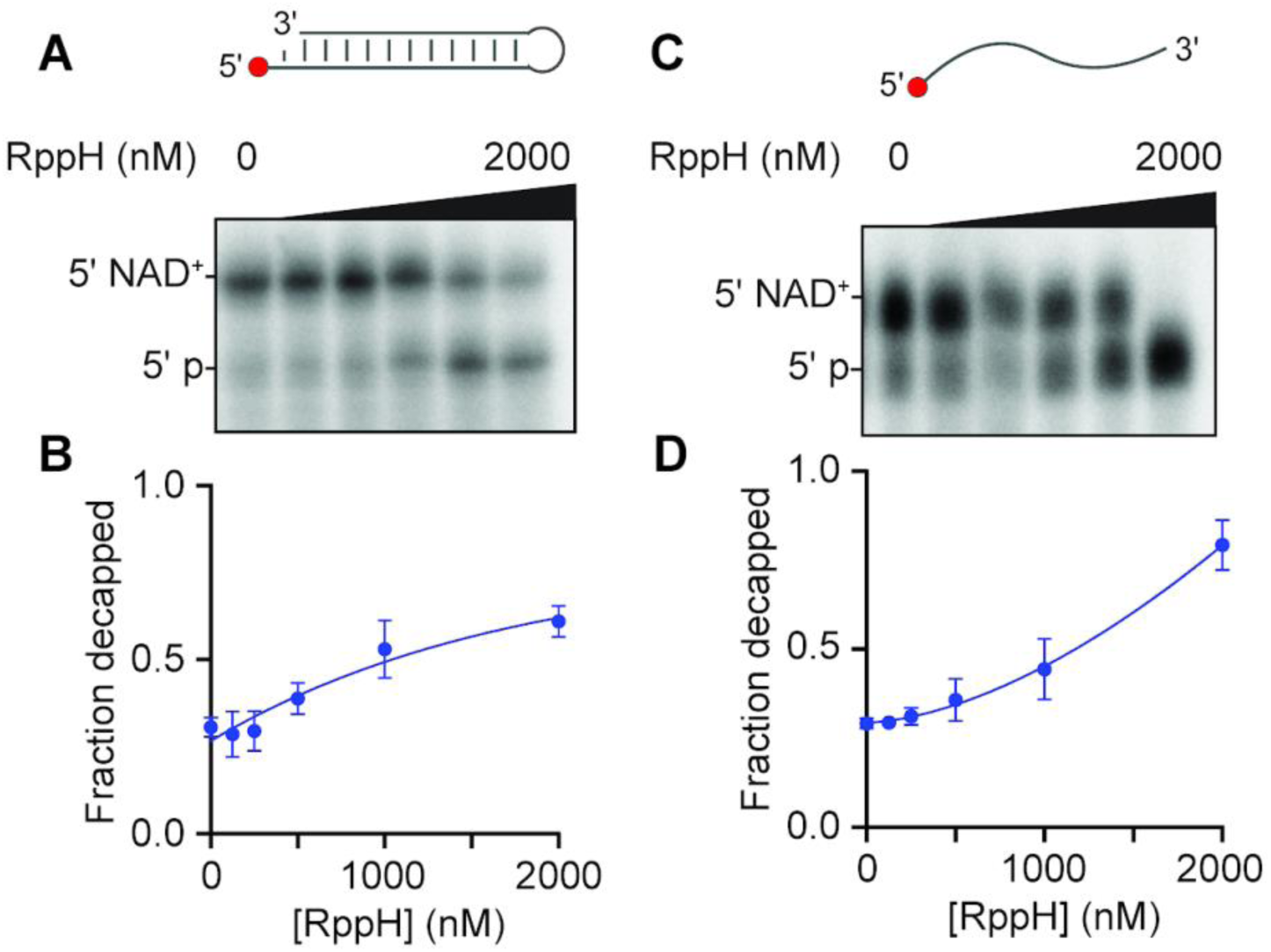
RppH decaps hairpin and linear NAD^+^-RNA substrates in vitro. (A) Representative urea-PAGE gel of concentration-dependent decapping of ³²P-labeled hairpin NAD^+^-RNA by RppH. RNA was incubated with 0–2 µM purified RppH for 30 min. (B) Quantification of decapping activity from the gel shown in A across three independent experiments. (C) Representative urea-PAGE gel showing concentration-dependent decapping of ³²P-labeled linear NAD^+^-RNA by RppH under the same conditions. (D) Quantification of decapping activity from the gel shown in C across three independent experiments. Data represent mean ± S.D. from three independent experiments (n = 3).

### GDP and ppGpp inhibit NAD(H)-RNA decapping by NudC and RppH

The ability of RppH and NudC to act on diverse cap structures led us to question whether free nucleotide species competitively inhibit decapping activity. To test inhibition, we monitored the ability of NudC to decap a hairpin NAD^+^-RNA substrate with a range of concentrations of ATP, GDP, GTP, or the nucleotide alarmone ppGpp (Fig. 2). While GTP and ATP had little effect on decapping, both GDP and ppGpp caused dose-dependent inhibition of NudC decapping activity (Fig. 2A-B; Supplementary Fig. S4) with IC_50_ values of ∼500 nM and ∼1 µM, respectively (Fig. 2). GDP levels in actively growing *E. coli* are reported to be ∼800 µM, which suggests that GDP must constitutively exert some level of inhibition on decapping during normal growth (27). The inability of ATP or GTP to inhibit NudC suggests that both the guanine base and the diphosphate moiety, which are present in both GDP and ppGpp, might be critical for inhibitory interactions with NudC. NudC was also able to efficiently decap the dinucleotide NAD^+^-RNA substrate in a concentration-dependent manner, and ppGpp potently inhibited this activity, with an IC_50_ of approximately 1.3 µM, demonstrating that ppGpp-mediated regulation extends to short NAD^+^-capped RNA substrates (Fig. 3). Amongst the longer RNA substrates, NudC has higher decapping activity toward NADH-RNA than NAD^+^-RNA *in vitro* (Fig S1). To determine whether ppGpp-mediated inhibition depends on the oxidation state of the NAD cap, we examined NudC-dependent decapping of NADH-capped RNA at increasing ppGpp concentrations. ppGpp inhibited decapping of NADH-capped RNA similarly to NAD^+^-capped RNA (Fig. 2D–E), indicating that ppGpp-mediated inhibition is independent of the oxidation state of the NAD cap.

**Figure 2.**
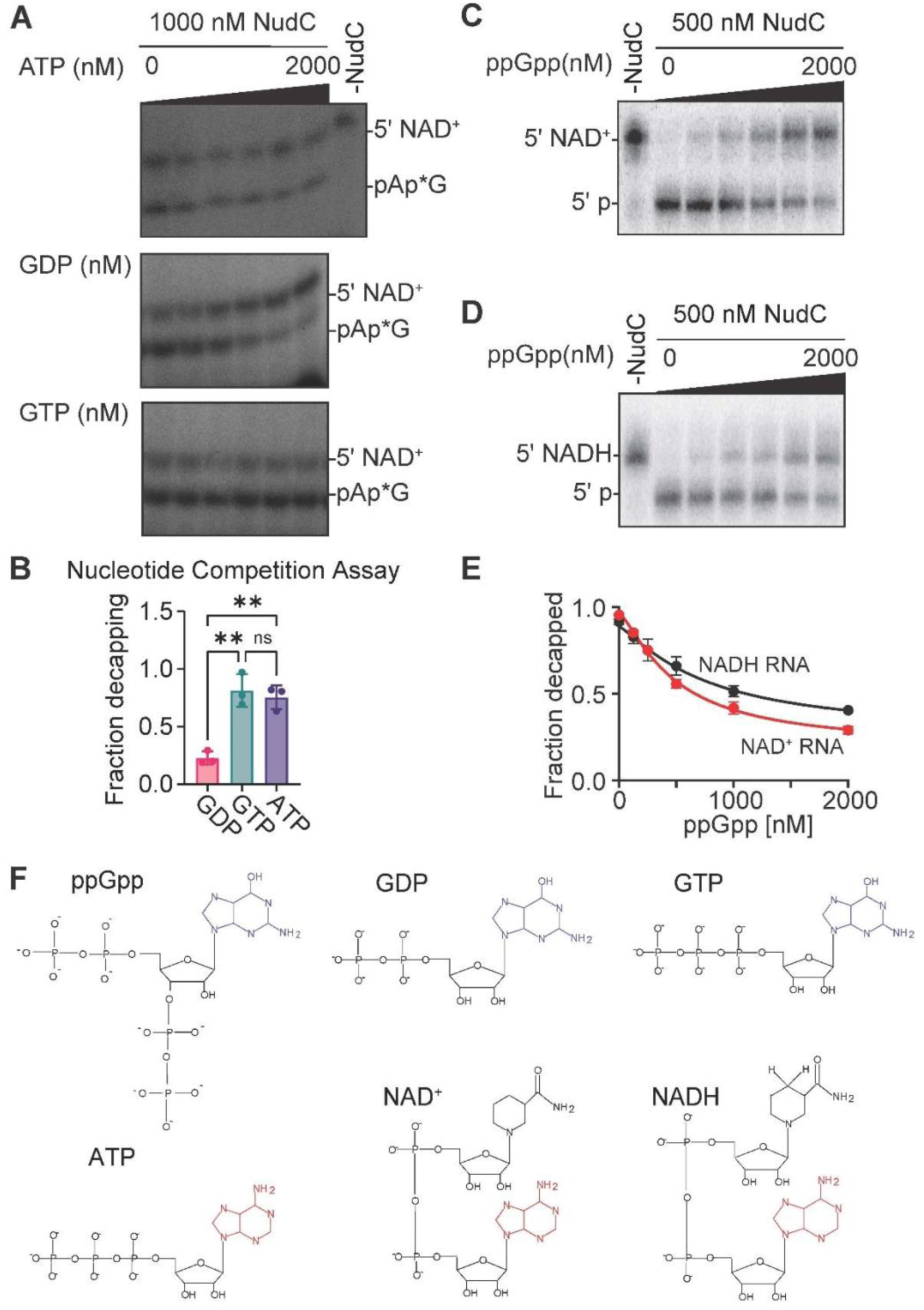
Regulation of NudC-mediated NAD(H)-RNA decapping by NTPs and NDP, and ppGpp on NAD(H)-RNA. (A) Representative decapping assays showing NudC activity on NAD-RNA in the presence of ATP, GTP, and GDP compared to no-NudC control. (B) Quantification of IC_50_ values for ATP, GTP, and GDP inhibition of NudC decapping activity as shown in A. (C) Decapping assay showing ppGpp-mediated inhibition of NudC activity on NAD+-RNA. (D) Decapping assay showing ppGpp-mediated inhibition of NudC activity on NADH-RNA. (E) Comparative quantification of ppGpp effects on NudC decapping of NAD RNA versus NADH-RNA. (F) Structural comparison of ppGpp, GDP, GTP, ATP, NAD^+^, and NADH; the adenine moiety shared among these molecules is highlighted in red to illustrate their common structural feature and its potential relevance to nucleotide recognition by NudC.

**Figure 3.**
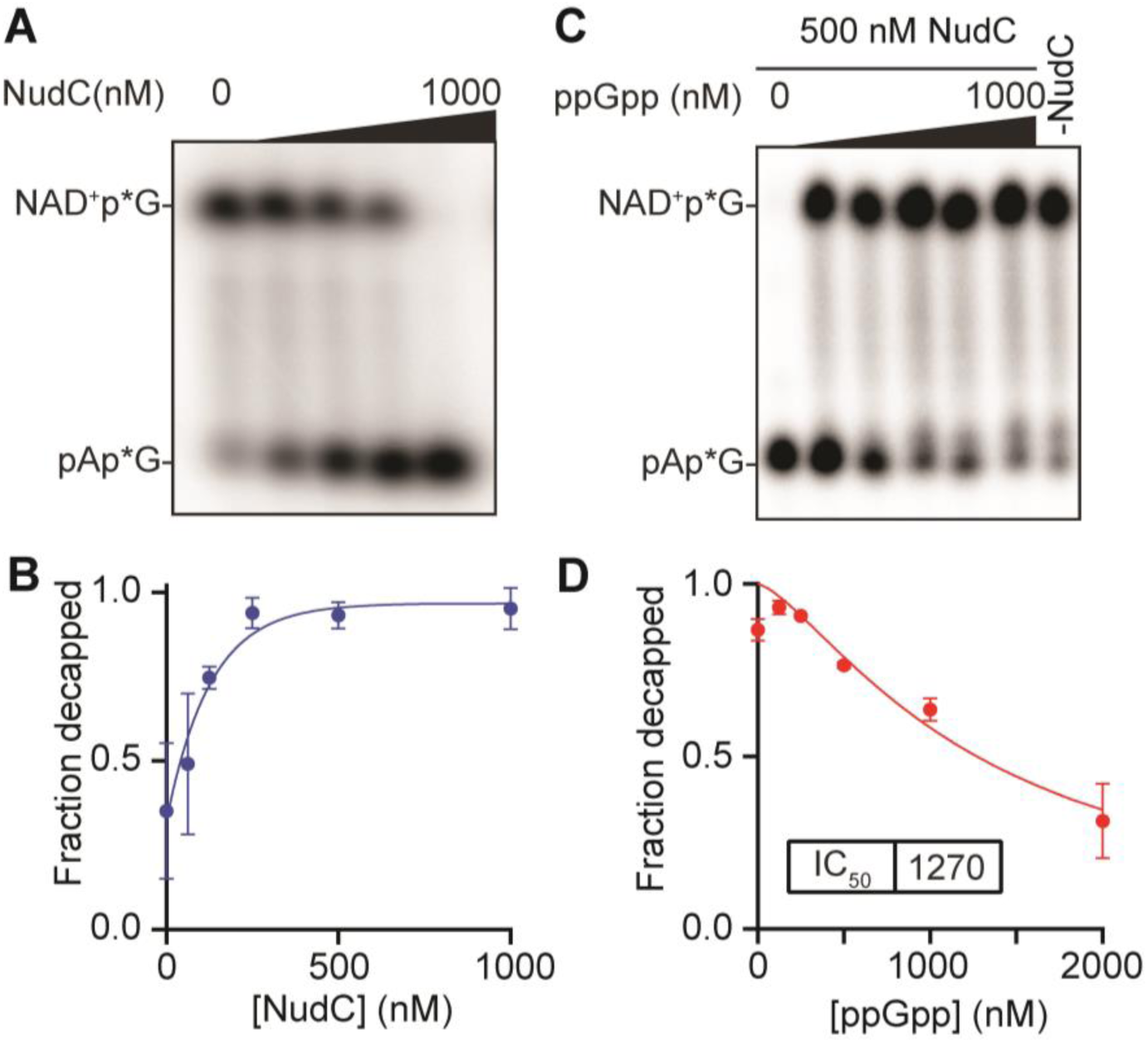
ppGpp inhibits NudC-mediated decapping of dinucleotide NAD^+^-RNA substrates. Urea-PAGE analysis of NudC decapping activity. (A) NudC dose-response assay (0–1000 nM) showing concentration-dependent decapping of short NAD^+^-RNA dinucleotides. (B) Decapping of dinucleotide NAD^+^-RNA by NudC (500 nM) in the presence of increasing concentrations of ppGpp (0–2000 nM), showing concentration-dependent inhibition. Data represent mean ± S.D. from three independent experiments (n = 3).

To determine whether ppGpp-mediated inhibition extends beyond NudC, we examined its effect on RppH-mediated decapping of NAD^+^-RNA. ppGpp inhibited decapping of both hairpin and linear NAD^+^-RNA substrates in a concentration-dependent manner (Fig. 4), with IC_50_ values of ∼500 nM and ∼600 nM, respectively. These values are comparable to the IC_50_ observed for inhibition NudC on hairpin RNA, indicating that both NAD-RNA decapping enzymes are similarly sensitive to ppGpp. Interestingly, ppGpp levels are known to fluctuate over a 100-fold range, from ∼10 µM during normal growth to ∼1 mM during nutrient starvation or stress (28, 29). Given the opportunity for dynamic changes in ppGpp levels to influence NAD^+^-RNA decapping, we sought to further characterize the inhibitory mode of action of this alarmone. Collectively, these results establish that ppGpp inhibits NAD^+^-RNA decapping by both NudC and RppH across dinucleotide and longer RNA substrates, with IC_50_ values ranging from approximately 0.5 to 1.3 µM (Table 1).

**Figure 4.**
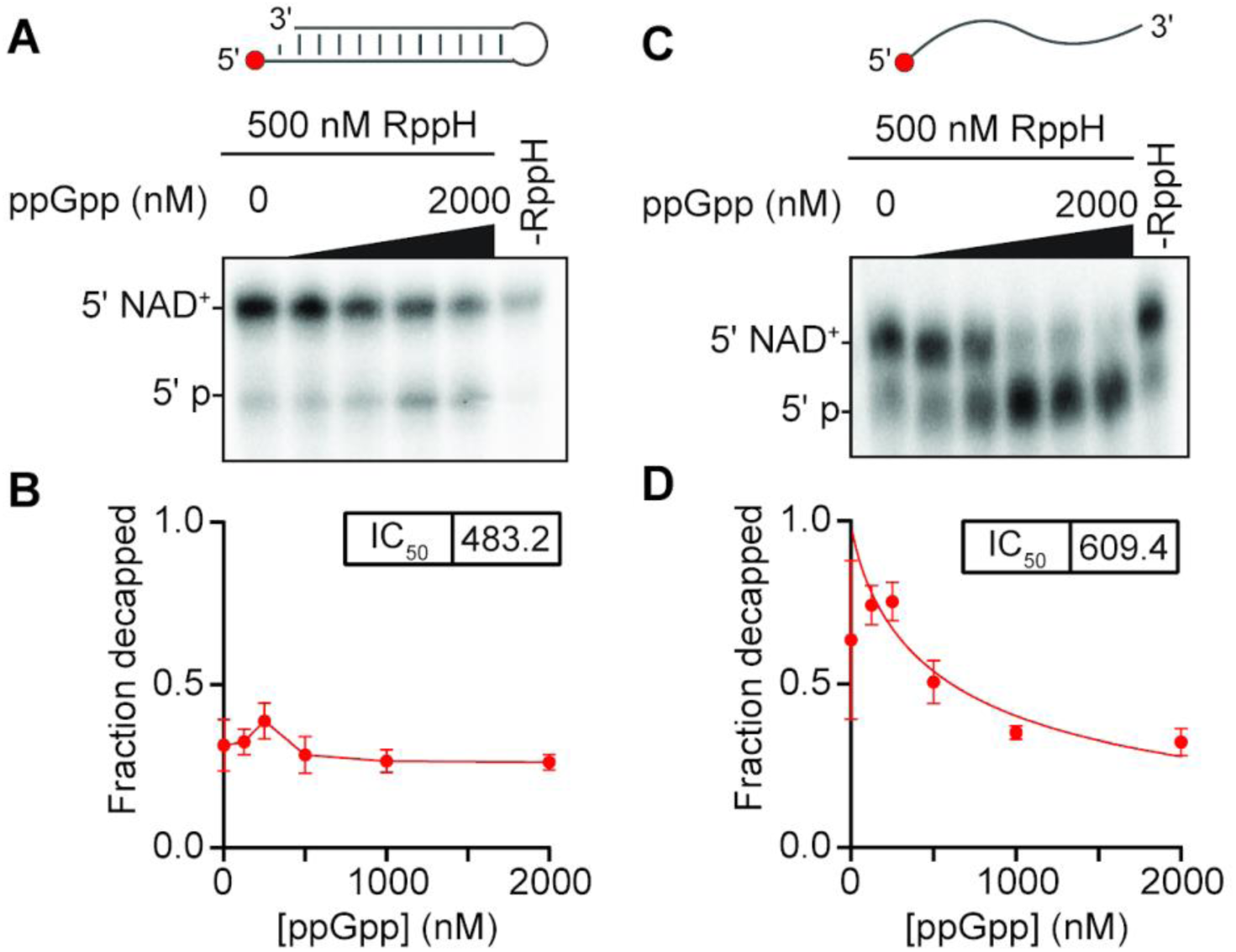
ppGpp inhibits RppH-mediated decapping of NAD^+^-RNA. Urea-PAGE analysis of ppGpp inhibition of RppH-mediated decapping using a fixed concentration of RppH (500 nM) and increasing concentrations of ppGpp (0–2000 nM). (A,B) Representative urea-PAGE gel and quantification showing ppGpp-dependent inhibition of RppH-mediated decapping of the hairpin NAD^+^-RNA substrate. (C, D) Representative urea-PAGE gel and quantification showing ppGpp-dependent inhibition of RppH-mediated decapping of the linear NAD^+^-RNA substrate. Data represent mean ± S.D. from three independent experiments (n = 3).

**Table 1:** IC_50_ values for ppGpp-mediated inhibition of NudC and RppH decapping of NAD^+^-capped RNA substrates.

| **Enzyme** | **NAD<sup>+</sup> capped RNA Substrate** | **IC<sub>50</sub> of ppGpp (nM)** |
|---|---|---|
| NudC | Dinucleotide RNA | 1270 |
| NudC | Hairpin Long RNA | 564 |
| RppH | Hairpin Long RNA | 483 |
| RppH | Linear Long RNA | 609 |

### The structural basis of NudC substrate specificity

NudC was originally identified as an NADH pyrophosphatase and has been shown to hydrolyze free NADH more efficiently than free NAD^+^. Subsequent studies demonstrated that NudC can also remove a variety of non-canonical RNA caps, including NAD-RNA and CoA-RNA, indicating broad substrate specificity toward metabolite-capped RNAs(4, 18, 19, 21, 30). To compare the relative susceptibility of NAD^+^-RNA and NADH-RNA to NudC-mediated decapping, we performed concentration-dependent dose-response assays using identical reaction conditions. NudC efficiently decapped both substrates; however, NADH-RNA was hydrolyzed at substantially lower enzyme concentrations than NAD-RNA (Fig. S1). Quantification of decapping activity reveals an approximately 10-fold increase in apparent activity toward NADH-RNA relative to NAD^+^-RNA. These results demonstrate a clear preference of *E. coli* NudC for NADH-capped RNA and are consistent with the previously reported preference of NudC for free NADH over free NAD^+^ (21). The enhanced activity toward NADH-RNA suggests that the redox state of the cap can influence substrate recognition and decapping efficiency.

To investigate the structural basis of NudC substrate recognition, we determined the crystal structure of *E. coli* NudC in complex with the reduced form of NADH at 1.55 Å resolution (Fig. 5A). (Crystal data collection and refinement statistics are provided in Table S1.) Clear positive *F_o_–F_c_* electron density was observed in the catalytic pocket, allowing unambiguous placement of the bound NADH molecule. The NudC dimer binds one NADH molecule in each active site, with NADH adopting an extended conformation (Fig. 5C). The adenine ring is stabilized by π-stacking with Y124 and F160, while the ribose hydroxyl groups form hydrogen bonds with Y124 and E111 (Fig. 5C).

**Figure 5.**
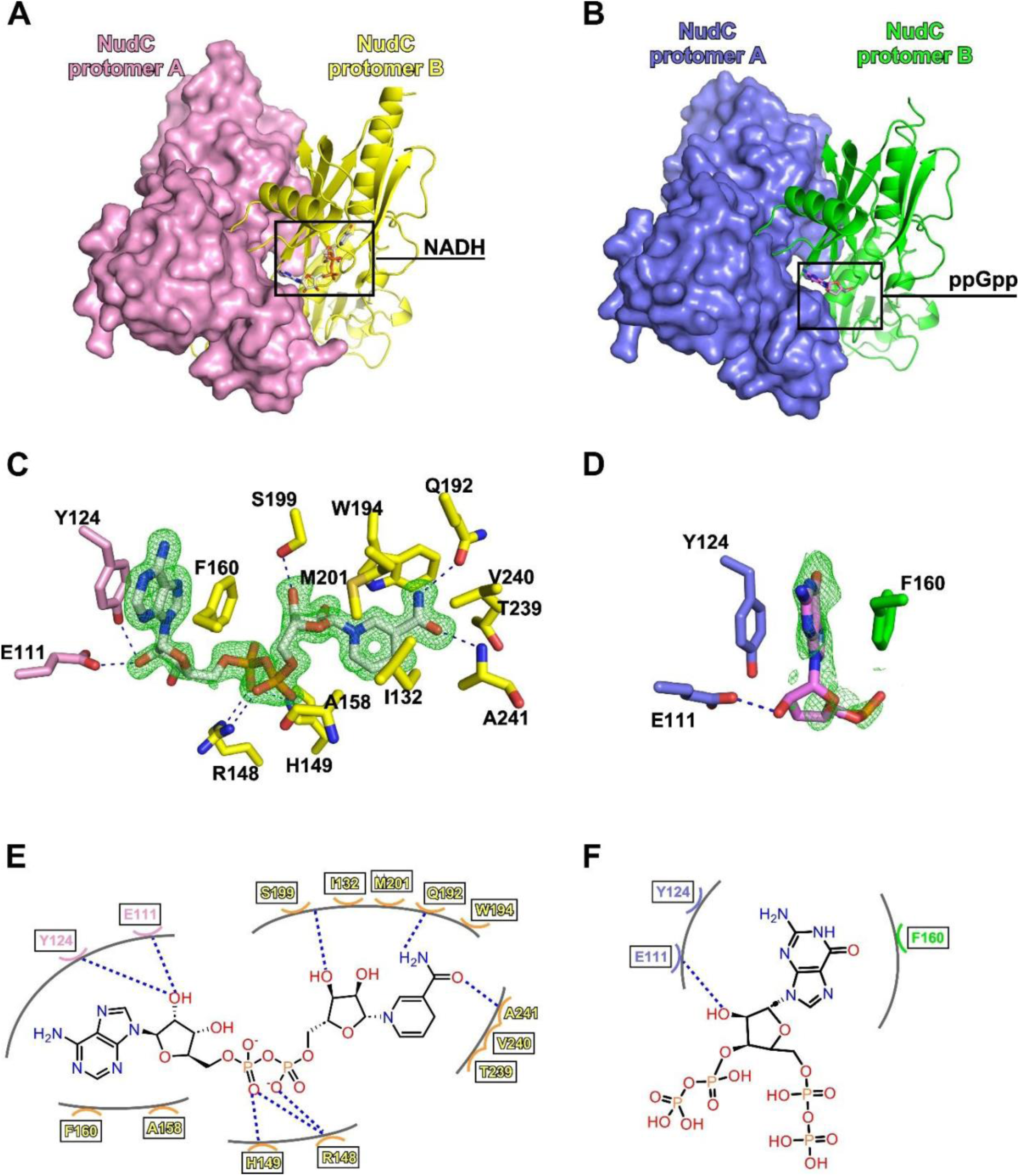
Structures and detailed interactions of *Ec*NudC in complex with NADH or ppGpp. (A) Overall structure of the *Ec*NudC–-NADH complex. The pink surface and yellow ribbon indicate individual protomers of *Ec*NudC. NADH is shown as white sticks. (B) Overall structure of the *Ec*NudC–-ppGpp complex. The light blue surface and green ribbon indicate individual protomers of *Ec*NudC. ppGpp is shown as magenta sticks. (C) Detailed interactions between *Ec*NudC and NADH. Pink sticks represent residues from protomer A of *Ec*NudC; yellow sticks represent carbon atoms of residues from protomer B; white sticks, carbon atoms of NADH; Nitrogen, oxygen, and phosphorus atoms are colored blue, red, and orange, respectively. (D) Detailed interactions between *Ec*NudC and ppGpp. Light blue sticks represent carbon atoms of residues from protomer A of *Ec*NudC; green sticks represent carbon atoms of residues from protomer B. (E) Schematic representation of the *Ec*NudC–NADH interactions. Boxes highlight residues that interact with NADH via side-chain atoms. (F) Schematic representation of the *Ec*NudC–ppGpp interactions. Boxes highlight residues that interact with ppGpp via side-chain atoms.

Previously, structures of NudC in complex with the oxidized form of NAD^+^ have been determined (18, 20). Notably, in our structures, the reduced nicotinamide moiety forms additional hydrogen-bonding and van der Waals interactions with the reduced NADH versus the oxidized NAD^+^ substrate used in previous structures: 1) Amino acids Q192 and A241 both form hydrogen bonds with the carboxamide side chain of the nicotinamide ring. 2) Amino Acids H149 and R148 contact the phosphate groups of the NAD structure, presumably holding the phosphates in the proper conformation to facilitate hydrolysis of the molecule. (Fig. 5E). Comparison of the NADH and NAD^+^-bound NudC structures revealed additional interactions in the NADH-bound complex that may contribute to substrate recognition and stabilization within the active site. These additional contacts provide a possible structural explanation for the enhanced decapping activity of NudC toward NADH-capped compared with NAD^+^-capped RNA.

### The structural basis of NudC inhibition by ppGpp

We next determined the crystal structure of NudC bound to ppGpp at 1.9 Å resolution (Fig. 5B; Table S1), revealing that ppGpp occupies the substrate-binding pocket within the NudC active site (Fig. 5D). The guanosine base is positioned between Y124 and F160, while E111 forms a hydrogen bond with the ribose hydroxyl group (Fig. 5F). The observed binding mode of ppGpp overlaps with the adenosine-binding region observed in the previously published NudC–NAD^+^ structure and our newly determined NudC–NADH structure, suggesting that ppGpp competes with NAD(H) for binding within the NudC active site and may hinder substrate access. Together with our biochemical data, these structures support a competitive inhibition mechanism for ppGpp-mediated inhibition of NudC.

Collectively, our findings identify RppH as an additional NAD^+^-RNA decapping enzyme in *E. coli* and demonstrate that ppGpp inhibits NAD^+^-RNA decapping by both RppH and NudC (Fig. 6). The structural data further supports a competitive mechanism of ppGpp-mediated inhibition of NudC, linking nucleotide signaling to the regulation of NAD^+^-capped RNA turnover.

**Figure 6.**
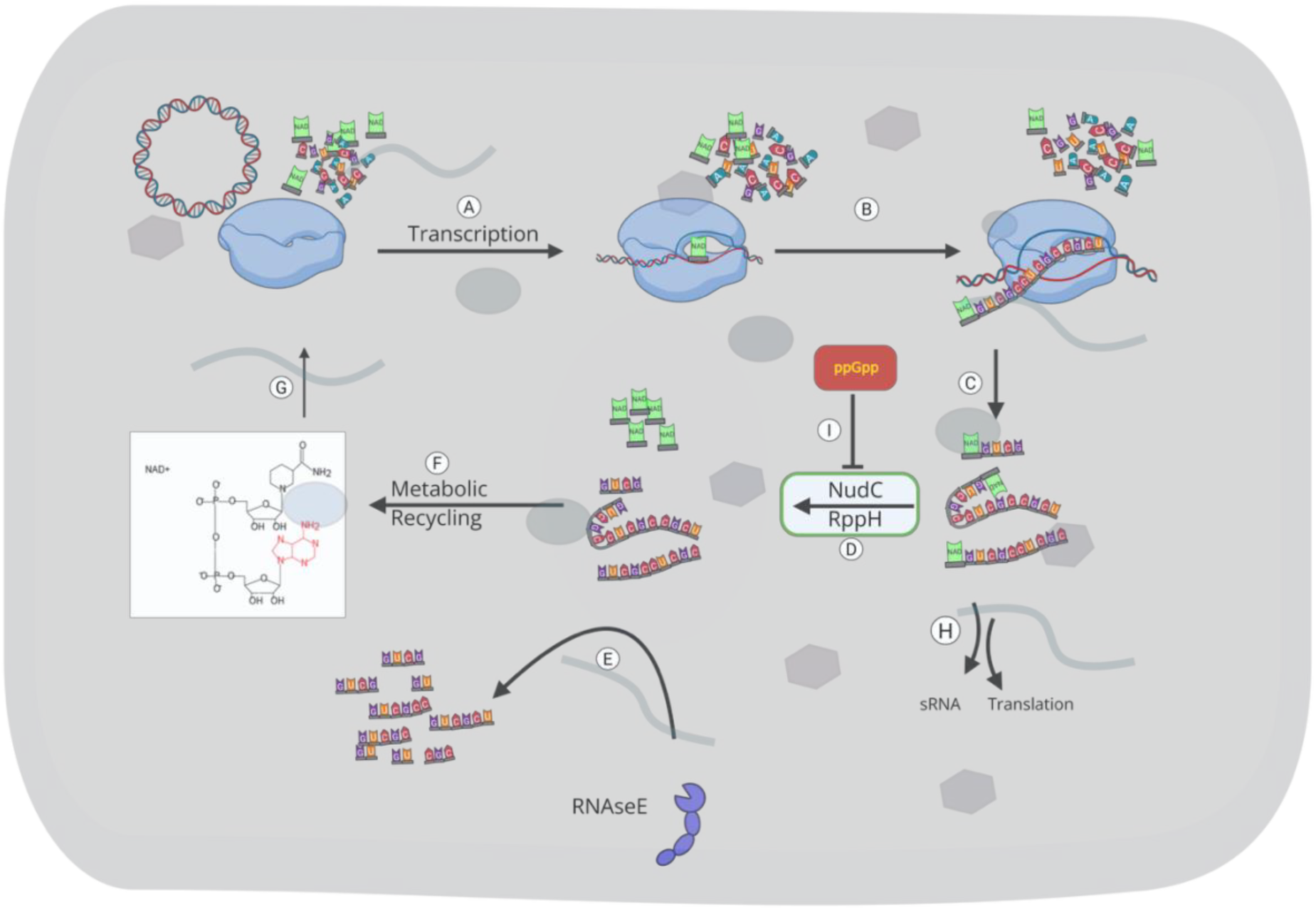
Graphical model linking NAD-capped RNA turnover to cellular metabolism and ppGpp signaling. NAD^+^ is incorporated at the 5’ end of RNA during transcription to generate NAD-capped RNA. The NAD cap can be removed by the decapping enzymes NudC and RppH, producing a 5’-monophosphorylated RNA that is subsequently targeted for RNase E–mediated degradation. Released NAD-derived metabolites may re-enter cellular metabolic pathways through recycling mechanisms. Accumulation of the alarmone ppGpp inhibits both NudC and RppH, thereby reducing decapping activity and promoting stabilization of NAD-capped RNAs during stress or stationary conditions.

## Discussion

Recent studies have solidified our understanding of how NAD^+^ caps are incorporated as non-canonical initiating nucleotides and which specific transcripts carry 5’-NAD^+^ caps (4, 11, 12, 30–32), but the functional consequences and regulatory dynamics of NAD^+^ capping remain less clear. NAD^+^-capped RNAs are protected from RNase E–mediated decay, and their turnover depends on decapping enzymes rather than the canonical 5’-monophosphate-dependent pathway(4, 18, 33). Until now, NudC was the only confirmed NAD-RNA decapping enzyme in *E. coli* (4, 18, 20). Here, we show that RppH also removes NAD^+^ caps from both linear and structured RNA substrates. Although, the observed activity of RppH on NAD^+^-capped RNAs was lower than that of NudC under comparable conditions (Fig. 1, S1). Even so, our data indicates that NAD-RNA turnover is governed by at least two Nudix hydrolases with distinct substrate preferences: NudC acting on both short and long NAD-RNAs, and RppH favoring longer substrates with an accessible 5’ end. This division of labor suggests that the two enzymes act on overlapping but non-identical pools of the NAD-RNA transcriptome, potentially providing some level of substrate-specific control over which capped transcripts are degraded.

In addition to identifying a second decapping enzyme, we also investigated the ability of ppGpp and other nucleotides to inhibit these decapping enzymes. Among the nucleotides we tested, only guanine-diphosphate-containing species (GDP and ppGpp) were inhibitory. This inhibition can be explained by the NudC-ppGpp crystal structure in which ppGpp occupies the same pocket used by the adenosine/nicotinamide moiety of NAD(H), with its guanine base sandwiched between residues Y124 and F160. These bases stack with the RNA substrate cap. Inhibition therefore arises from steric competition at the active site, rather than allosteric regulation. This may explain why the functionally similar NudC and RppH are both sensitive to ppGpp despite limited sequence identity. While GDP and ppGpp are similarly potent inhibitors *in vitro*, ppGpp is the more plausible mediator of dynamic decapping regulation because its concentration changes by roughly two orders of magnitude between exponential growth and stress, whereas GDP is present at comparatively stable levels throughout growth. The similar IC_50_ values we observed for ppGpp across NudC and RppH, and across dinucleotide, hairpin, and linear substrates, argue that elevated ppGpp levels during stress would inhibit both enzymes, applying a coordinated brake on NAD-RNA turnover precisely when NAD-RNA levels are known to increase. Although NudC and RppH share a conserved Nudix catalytic motif, structural comparisons suggest that their substrate-binding pockets differ in orientation and organization, potentially contributing to their distinct substrate preferences (Supplementary Fig. S3).

Our findings support the view that metabolite-capped RNAs are actively regulated species rather than passive by-products of transcription. Alongside prior reports that ppGpp stimulates NAD^+^ incorporation on transcripts such as *sibD* during transcription initiation (34), our work suggests that ppGpp stimulates stress-induced NAD-RNA accumulation through a two-pronged mechanism, by simultaneously promoting NAD-RNA synthesis and blocking NAD^+^ removal. This coordinated influence on synthesis and decay by a single metabolic signal would produce a more robust shift in the NAD-capped transcriptome than regulating transcription alone. This regulatory paradigm is consistent with ppGpp’s broader role in regulating RNA metabolism during the stringent response, including its inhibition of RppH-mediated RNA pyrophosphohydrolysis and PNPase-mediated RNA degradation in *Streptomyces* species(17, 25).

Several questions follow from this model. It remains unknown which endogenous transcripts are the physiological substrates of RppH-mediated NAD-decapping *in vivo*, and whether the NudC/RppH division of labor we observe biochemically holds across the cellular NAD-RNA pool. It is also unclear whether ppGpp regulates additional Nudix family members beyond these two, which would suggest a more general mechanism for coupling alarmone signaling to RNA cap metabolism. Future studies must address these questions to establish the physiological significance of the regulatory mechanism we describe here.

## Materials and Methods

### Plasmid construction, strain, and purification of NudC for functional activity analysis

*E. coli* NudC protein was purified as described in Cahova *et al.* (2015; 4). The pET28c-*EcnudC* plasmid encoding *nudC* with a C-terminal 6×His tag in the pET28c kanamycin-resistant expression vector was transformed into NiCo21(DE3) competent *E. coli* cells (New England Biolabs). For protein expression, a starter culture was used to inoculate LB medium containing kanamycin and grown at 37°C with shaking until the culture reached an OD_600_ of ∼0.6. Expression of His-tagged NudC was induced with 1 mM IPTG, and incubation was continued for 2 hours at 37°C. Cells were harvested by centrifugation and resuspended in lysis buffer (Tris 50mM, NaCl 1 M, urea 1M, MgSO_4_ 5mM, imidazole 10mM, glycerol 5%, lysozyme 3mg/mL, protease inhibitor (EDTA-free) - 1 tablet per 100mL). Cells were lysed by sonication, and insoluble material was removed by centrifugation. The clarified lysate was incubated with Ni²^+^-NTA affinity resin (Molecular Cloning Laboratories) for 1 hour at 4°C with gentle mixing. Bound protein was washed with wash buffer containing imidazole (30 mM), and NudC was eluted using elution buffer containing 200 mM imidazole. Eluted fractions containing NudC were spin-concentrated and further purified by size-exclusion chromatography on a Superdex 75 16/60 column (Cytiva) using an ÄKTA FPLC system equilibrated in storage buffer (50 mM Tris pH 8.0, 1M NaCl, 2 mM DTT, 5% Glycerol). Peak fractions were pooled and concentrated using Amicon Ultra centrifugal filters (10 kDa MWCO, Millipore Sigma). Protein purity was assessed by SDS-PAGE (Fig. S2). Purified NudC was aliquoted and stored in storage buffer supplemented with 50% glycerol at –80°C.

### NudC protein purification for crystallography

For the preparation of *Escherichia coli* NudC (*Ec*NudC) for crystallography, *E. coli* BL21(DE3) cells harboring the pET28c-*EcnudC* plasmid were cultivated in Luria-Bertani (LB) medium at 37°C. When the optical density at 600 nm (OD_600_) reached 0.6, the expression of C-terminal 6×His-tagged *Ec*NudC was induced with 0.5 mM isopropyl β-D-1-thiogalactopyranoside (IPTG) at 16°C for 14 h. Cells were harvested by centrifugation at 8000 × g and 4°C, then resuspended in lysis buffer (50 mM Tris-HCl, pH 8.0, 1 M NaCl, 5 mM imidazole, 5 mM MgSO_4_, 5% glycerol, 2 mM phenylmethylsulfonyl fluoride (PMSF), 5 mM β-mercaptoethanol (β-ME). Cell lysis was performed using an Avestin EmulsiFlex-C3 cell disrupter (Avestin, Inc.). The supernatant was collected by centrifugation at 16,000 × g for 45 min at 4°C and loaded onto a 2 mL Ni-NTA agarose column (SMART-Lifesciences). The column was washed with lysis buffer supplemented with 20 mM imidazole, and the bound protein was eluted with lysis buffer containing 150 mM imidazole. The eluted fraction was dialyzed overnight against dialysis buffer (10 mM Tris-HCl, pH 8.0, 50 mM NaCl, 1 mM dithiothreitol (DTT)).

The dialyzed sample was further purified using a HiTrap Q HP anion-exchange column (Cytiva, Inc.) with a linear gradient composed of buffer A (10 mM Tris-HCl, pH 8.0, 50 mM NaCl, 1 mM DTT) and buffer B (10 mM Tris-HCl, pH 8.0, 1 M NaCl, 1 mM DTT). Fractions containing *Ec*NudC were concentrated to 2 mL and loaded onto a HiLoad 16/60 Superdex 75 column (Cytiva, Inc.) pre-equilibrated with 10 mM Tris-HCl, pH 8.0, 50 mM NaCl, 1 mM DTT. The target protein fractions were collected, concentrated to 10 mg/mL, and stored at –80°C for further use.

### Structure determination of EcNudC-NADH

*Ec*NudC at a concentration of 10 mg/mL was incubated with 2 mM NADH for 30 minutes, after which initial crystallization screens were performed using the sitting-drop vapor diffusion method. Diffraction-quality crystals were harvested following 5 days of growth at 22°C in optimized reservoir solution A (0.1M Bis-Tris pH 5.5, 15% PEG3350, 0.2M sodium malonate). Prior to data collection, crystals were soaked in 10% (v/v) (2R, 3R)-(-)-2,3-butanediol as a cryoprotectant and flash-cooled in liquid nitrogen. X-ray diffraction data were collected at beamline 17U of the Shanghai Synchrotron Radiation Facility (SSRF) and processed with the HKL2000 software suite. The structure was solved by molecular replacement using Phaser MR, with the structure of *Ec*NudC in complex with NADH (PDB ID: 5ISY) employed as the search model. The resulting electron density maps exhibited clear and well-defined density for both *Ec*NudC and the bound NADH ligand. Iterative rounds of model building and structure refinement were subsequently conducted using Coot and Phenix, respectively. The final structure of the *Ec*NudC–NADH complex has been deposited in the Protein Data Bank with the accession code 24MG.

### Structure determination of EcNudC-ppGpp

*Ec*NudC at a concentration of 10 mg/mL was incubated with 2 mM ppGpp for 30 min, after which initial crystallization screens were performed using the sitting-drop vapor diffusion method. Diffraction-quality crystals were harvested following 8 days of growth at 22°C in optimized reservoir solution B (1% w/v Tryptone, 0.05 M HEPES sodium, pH 7.0, 12% w/v polyethylene glycol 3,350). Prior to data collection, crystals were soaked in 10% (v/v) (2R, 3R)-(-)-2,3-butanediol as a cryoprotectant and flash-cooled in liquid nitrogen. X-ray diffraction data were collected at beamline 17U of the Shanghai Synchrotron Radiation Facility (SSRF) and processed with the HKL2000 software suite. The structure was solved by molecular replacement using Phaser MR, with the structure of Apo *Ec*NudC in complex with NADH employed as the search model. The resulting electron density maps exhibited well-defined density for *Ec*NudC. In contrast, the electron density for ppGpp was less well-defined, such that only the guanosine moiety could be modeled. Iterative rounds of model building and structure refinement were subsequently conducted using Coot and Phenix software, respectively (27, 28). The final structure of the *Ec*NudC–ppGpp complex has been deposited in the Protein Data Bank with the accession code 24MH.

### Preparation of linear and hairpin RNA transcription templates

To examine the decapping activity of NudC and RppH on substrates with different structural features, two RNA classes were used: a linear RNA lacking secondary structure near the 5’ end, and a hairpin RNA containing a defined secondary structure.

DNA transcription templates were generated by annealing complementary template and non-template oligonucleotides (Material and Method Table 1). Oligonucleotides (source IDT) were mixed at final concentrations of 2.0 µM template strand and 2.2 µM non-template strand in nuclease-free water. Samples were heated to 95°C for 10 minutes and slowly cooled to room temperature at a rate of 1°C per minute to allow annealing. Annealed ds templates were stored at –20°C until used for in vitro T7 transcription reactions.

### In-vitro transcription of NADpG dinucleotide RNA

NAD^+^-linked dinucleotide transcripts consisting of NAD^+^-p-RNA species were generated by in-vitro transcription as described previously (Singh and Bird, 2022) (29). Transcription reactions were carried out using T7 RNA polymerase (New England Biolabs) in 1× transcription buffer containing 40 mM Tris-HCl, 6 mM MgCl2, 1 mM DTT, and 2 mM spermidine. Final reactions (50 µL) additionally contained NAD^+^ (10 mM), GTP (20 µM; Cytiva, Inc.), DNA template (2 µM) (Table S1), and [α-³²P] GTP (0.5 µL per reaction; Revvity, Inc.) to radiolabel the transcript.

Reaction mixtures were incubated at 37°C for 30 minutes. Following transcription, products were purified using NanoSep centrifugal columns (Pall Nanosep columns, 10k) to remove protein, DNA template, and unincorporated nucleotides. Purified dinucleotide NAD-RNA products were used directly as substrates for decapping assays with NudC or RppH in the presence or absence of ppGpp.

### In vitro transcription of full-length NAD-capped RNA

Full-length NAD^+^-capped RNA substrates were generated by in-vitro transcription using T7 RNA polymerase and 2uM A-less templates (hairpin and linear) (Table S2) under similar conditions to those used for dinucleotide NAD-RNA synthesis. Transcription reactions contained NAD^+^ as the initiating nucleotide together with the extending nucleoside triphosphates GTP, UTP, and CTP, as well as [α-³²P] GTP to radiolabel the transcript to make a 50 µL reaction. Reactions were incubated at 37°C for 30 minutes. Following transcription, full-length RNA products were purified using Illustra MicroSpin G-25 columns (Cytiva) to remove unincorporated nucleotides and small-molecule reaction components. Columns were equilibrated according to the manufacturer’s instructions by centrifugation at 735 × g for 1 min to remove storage buffer. Transcription reactions were then applied to the column and centrifuged for 2 minutes to recover purified RNA. Purified RNA was ethanol precipitated in the presence of glycogen stop solution (glycogen 100ug/mL + 50mM of NaOAc Solution). After overnight incubation in -20°C, RNA was spun down (13,000 rpm, 4°C, 30 minutes), RNA pellets were air-dried briefly and resuspended in decapping reaction buffer 1x NEBuffer 2 (New England Biolabs; containing Tris-HCl, NaCl, and MgCl_2_) and incubated on ice for at least 15 minutes to allow complete dissolution before decapping assays. This method is provided in detail in Singh and Bird (2022; (35).

### Enzymatic decapping assays with NudC and RppH

Decapping reactions were performed in a final volume of 10 µL containing 9 µL radiolabeled capped RNA substrate in 1× NEBuffer and 1 µL purified NudC or RppH enzyme. Enzyme concentrations were varied as indicated, from 125 nM to 2000 nM final concentration Nudc and 2000nM RppH, by addition of 1 µL of appropriately diluted protein stock. Control reactions had no enzyme and 1 µL of protein storage buffer.

Reactions were incubated at 37°C for 30 minutes and terminated by the addition of an equal volume of transcription loading buffer (TLB) (1M Tris, pH 8.0, 20% SDS, 0.5M EDTA, deionized formamide). Reaction products were stored in -20°C until gel run. Reaction products from dinucleotide substrates were separated on 0.4mm thickness 20% denaturing polyacrylamide urea gels (19:1 Urea-gel National Diagnostics). Full-length RNA substrates were analyzed on 0.4mm thickness 10% denaturing polyacrylamide gels. 0.5% phenylboronic acid (PBA) (Boron Molecular, Inc.) is dissolved into the gel to improve the separation of NAD-capped and decapped RNA species (36).

Following electrophoresis, gels were transferred to Whatman paper, dried under vacuum, and exposed to phosphor imaging screens overnight (Cytiva, Inc.). Screens were scanned using an Azure Sapphire biomolecular imager (Azure Biosystems). Band intensities were quantified using ImageLab software (Bio-Rad).

### Inhibition assays of NudC and RppH by ppGpp

To evaluate inhibition of decapping activity by ppGpp, reactions were performed on dinucleotide, and both linear and structured RNA substrates under conditions identical to the standard decapping assays except that enzyme concentration was held constant, and ppGpp (stock 20uM) was titrated. For NudC inhibition assays, NudC was used at a concentration corresponding to near-complete substrate decapping under the assay conditions (500 nM final concentration). Each 10 µL reaction contained capped RNA substrate in 1× NEBuffer 2, 1 µL NudC, and 1 µL ppGpp (Axxora/Enzo Life Sciences) diluted to give final ppGpp concentrations ranging from 125 nM to 2000 nM. Control reactions lacking ppGpp or NudC received an equivalent volume of buffer. Reactions were incubated at 37°C for 30 minutes and terminated by the addition of an equal volume of TLB. Products were resolved by denaturing urea-PAGE as described above and visualized by phosphor imaging. Decapping activity in the presence of ppGpp was quantified relative to control reactions lacking the inhibitor, and inhibition curves were fitted using a nonlinear model to determine IC_50_ values.

Equivalent inhibition assays were performed for RppH using fixed enzyme concentrations and increasing ppGpp concentrations on both linear and structured RNA substrates under the same reaction conditions. Where required, enzyme storage buffer was added to maintain a constant final reaction volume of 10 µL.

### Inhibition assays of NudC by GTP, ATP, and GDP

To compare the specificity of NudC inhibition by related nucleotides, decapping assays were performed as described above using a fixed concentration of NudC (500nM) and increasing concentrations of GDP, GTP, or ATP in place of ppGpp. Nucleotides were tested over the same final concentration range used for ppGpp (125–2000 nM). Reactions were incubated at 37°C for 30 min, terminated with TLB, and analyzed by denaturing PAGE followed by phosphor imaging. Decapping activity was quantified relative to reactions lacking added nucleotide.

### Gel electrophoresis, imaging, and quantification

Reaction products from full-length RNA substrates were resolved on 10% denaturing polyacrylamide gels, whereas dinucleotide substrates were analyzed on 20% denaturing polyacrylamide gels containing urea (National Diagnostics). For the separation of NAD-capped and decapped full-length RNA species, 10% gels were supplemented with 0.5% PBA.

Gels were electrophoresed until free nucleotides migrated to the bottom of the gel (typically ∼1 hour at 35W constant power). 1x TBE was used as a running buffer. Following electrophoresis, gels were transferred to Whatman paper. Full-length RNA gels were vacuum-dried using a Bio-Rad Gel Dryer Model 583 before imaging. Dried gels were exposed to phosphor imaging screens overnight and scanned using an Azure Sapphire phosphor imager (Azure Biosystems).

Band intensities were quantified using Image Lab software (Bio-Rad). For decapping assays, the fraction decapped was calculated as the intensity (absolute volume) of the lower (decapped) band divided by the total lane signal (sum of capped and decapped species). Data from three independent experiments (n = 3) with error bars = SD were plotted and analyzed using GraphPad Prism.

## Acknowledgements

We would like to thank Dr. Karl Schmitz for helpful discussions and technical assistance;

Laura Marvin and Kaustuv Das for technical assistance; and Jared Schrader and Boyuan Wang for discussions.

## Funding

This work was supported by Startup funds provided by the University of Delaware College of Arts and Sciences.

## Author Contribution

Conceptualization: Jeremy Bird, Yu Zhang

Methodology: Yogeshwari Singh, Yu Zhang and Jeremy Bird

Formal Analysis: Yogeshwari Singh, Lingting Li, Yu Zhang and Jeremy Bird

Investigation: Yogeshwari Singh, Raymond George and Lingting Li

Writing: Yogeshwari Singh and Jeremy Bird

Visualization: Yogeshwari Singh, Lighting Li, Yu Zhang and Jeremy Bird

Supervision: Jeremy Bird and Yu Zhang

Project Administration: Jeremy Bird Funding Acquisition: Jeremy Bird.

## Conflict of Interest Statement

The authors declare no conflict of interest.

## Disclosure of AI use

ChatGPT (OpenAI) was used to assist with language editing, including improving sentence clarity, grammar, and readability.

## SUPPLEMENTARY DATA

**Supplementary Figure S1.**
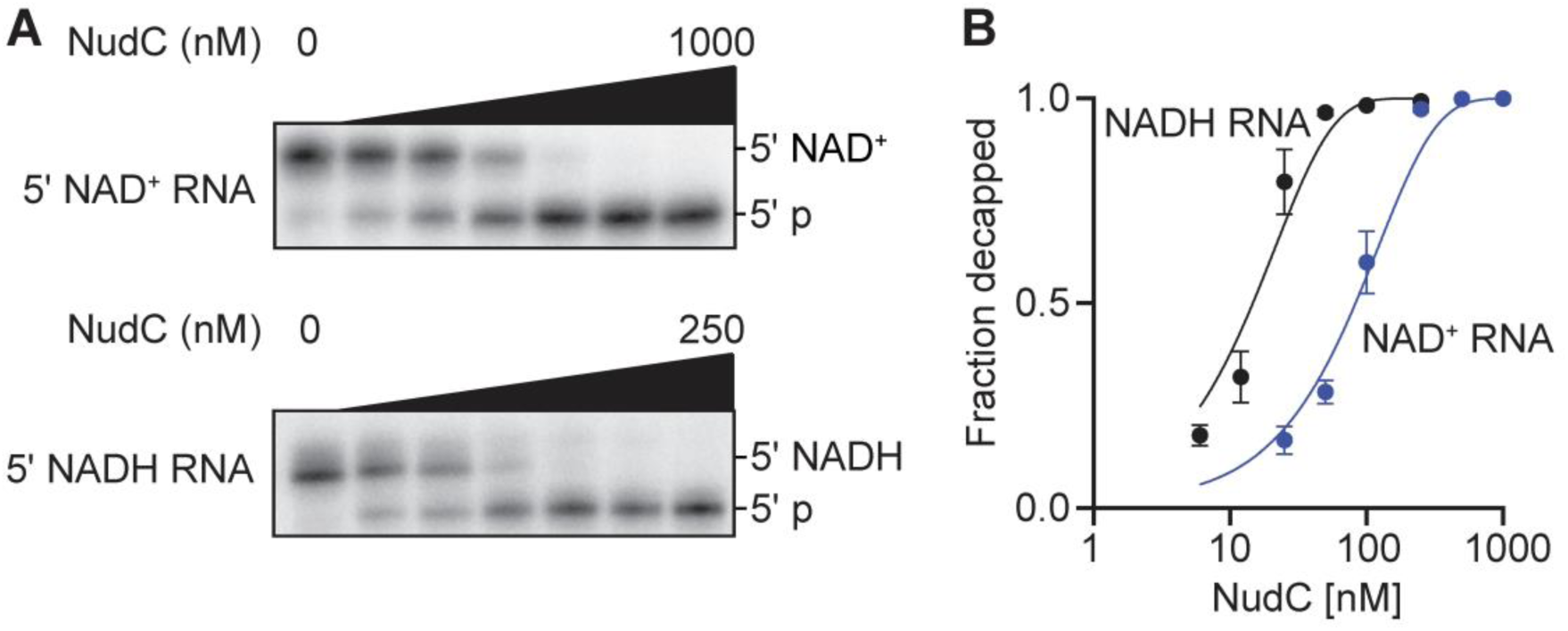
NudC preferentially decaps NADH-RNA over NAD⁺-RNA. Urea-PAGE analysis of NudC-mediated decapping of NAD⁺-RNA and NADH-RNA across increasing NudC concentrations under identical assay conditions. (A) Representative gels showing concentration-dependent conversion of NAD⁺ and NADH-capped RNA to decapped product. (B) Quantification of decapping efficiency as a function of NudC concentration. NudC exhibits higher activity toward NADH-RNA than NAD⁺-RNA, demonstrating substrate preference for the reduced nicotinamide cap. Data represent mean ± s.d. from independent experiments (n = 3).

**Supplementary Figure S2.**
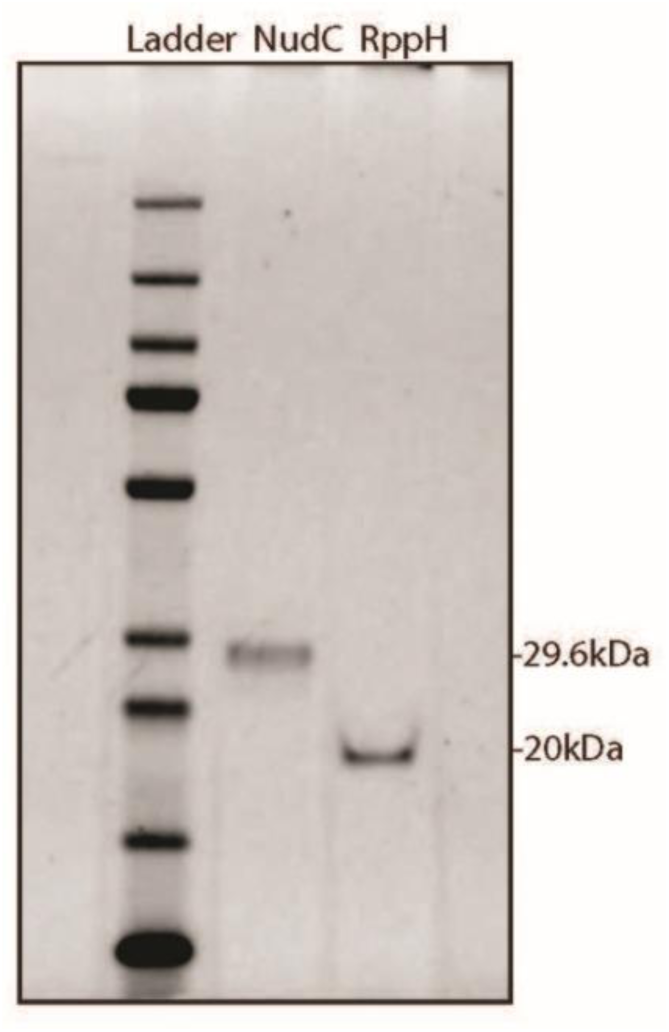
Purification and quality assessment of NudC and RppH proteins used in decapping assays-. SDS-PAGE analysis of proteins used throughout this study. (a) Purified His-tagged *E. coli* NudC (∼29.6 kDa) following nickel affinity and size-exclusion chromatography. (B) *E. coli* RppH (∼19 kDa) protein (NEB). Proteins were resolved by SDS-PAGE and visualized by Coomassie staining. The prominent bands at the expected molecular weights indicate high purity of both proteins used for biochemical and enzymatic assays.

**Supplementary Figure S3.**
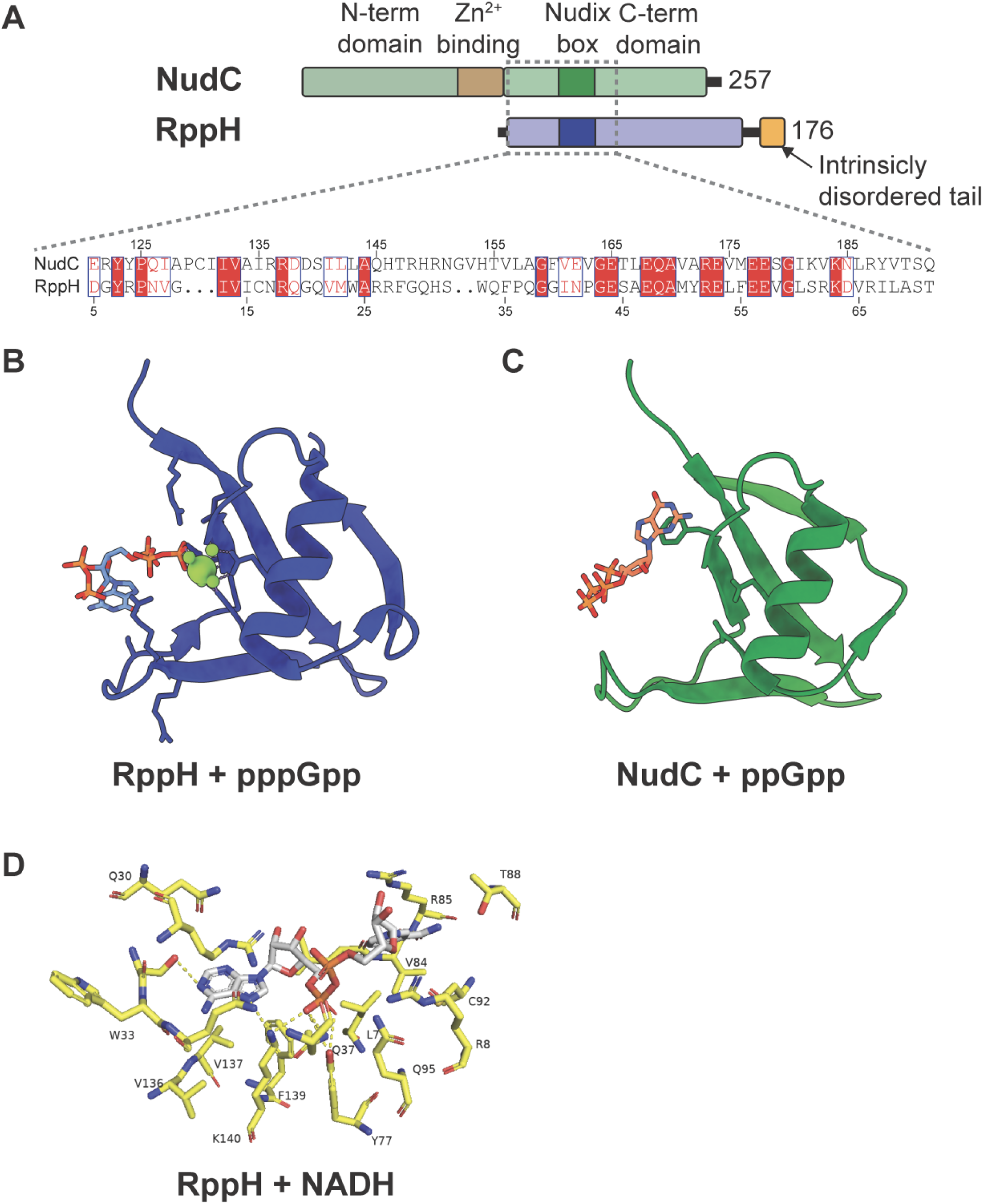
Structural comparison of *E. coli* RppH (blue) and NudC (green) reveals conservation of the Nudix catalytic motif -. (A) Sequence alignment of *E. coli* NudC and RppH, highlighting conserved residues within the Nudix domain including the nudix box motif. Conserved amino acids are block coloured red and similar amino acids are shown in red text (B) Model of pppGpp-bound RppH displayed as a cartoon representation to illustrate the potential positioning of inhibitor within the RppH active site relative to the Nudix domain (blue). Magnesium ions shown in blue and pppGpp in stick representation. Structural data from Gao et al. (2020); PBD: 6VCL. (C) Positioning of ppGpp in the active center of the NudC Nudix domain (blue) from the structures generated in this work (PDB: 24MH). Comparison of B and C demonstrate that both enzymes have differentially oriented substrate pockets despite both being inhibited by Guanosine containing nucleotides in a similar manner. (D) Model of NADH-RppH, showing NADH positioning in the substrate binding pocket generated using Protenix-v2(1). Yellow sticks represent amino acids of RppH and white sticks the NADH moiety in the binding site. Together, these analyses highlight conservation of the nudix catalytic scaffold while suggesting distinct substrate-binding features that may contribute to the different substrate preferences of NudC and RppH.

**Supplementary Figure S4.**
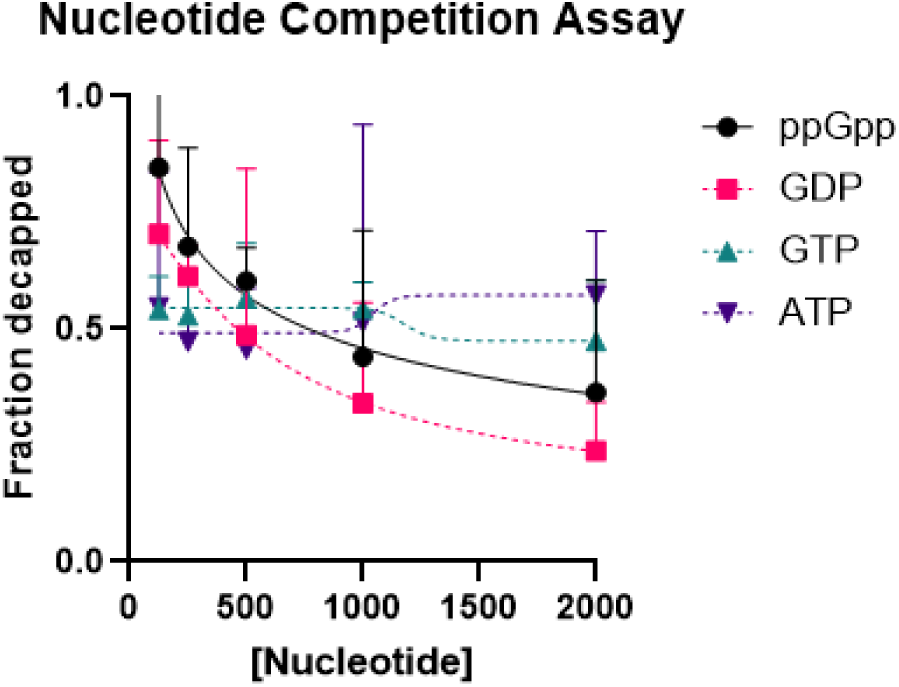
Selective inhibition of NudC-mediated NAD-RNA decapping by guanine diphosphate-containing nucleotides-. Competition assay comparing the effects of ATP, GTP, GDP, and ppGpp on NudC-mediated decapping of NAD⁺-capped RNA. NudC activity was measured in the presence of increasing concentrations of each nucleotide and quantified as the fraction of RNA decapped. Data points represent mean ± SD (n = 3).

**Supplementary Table 1.**
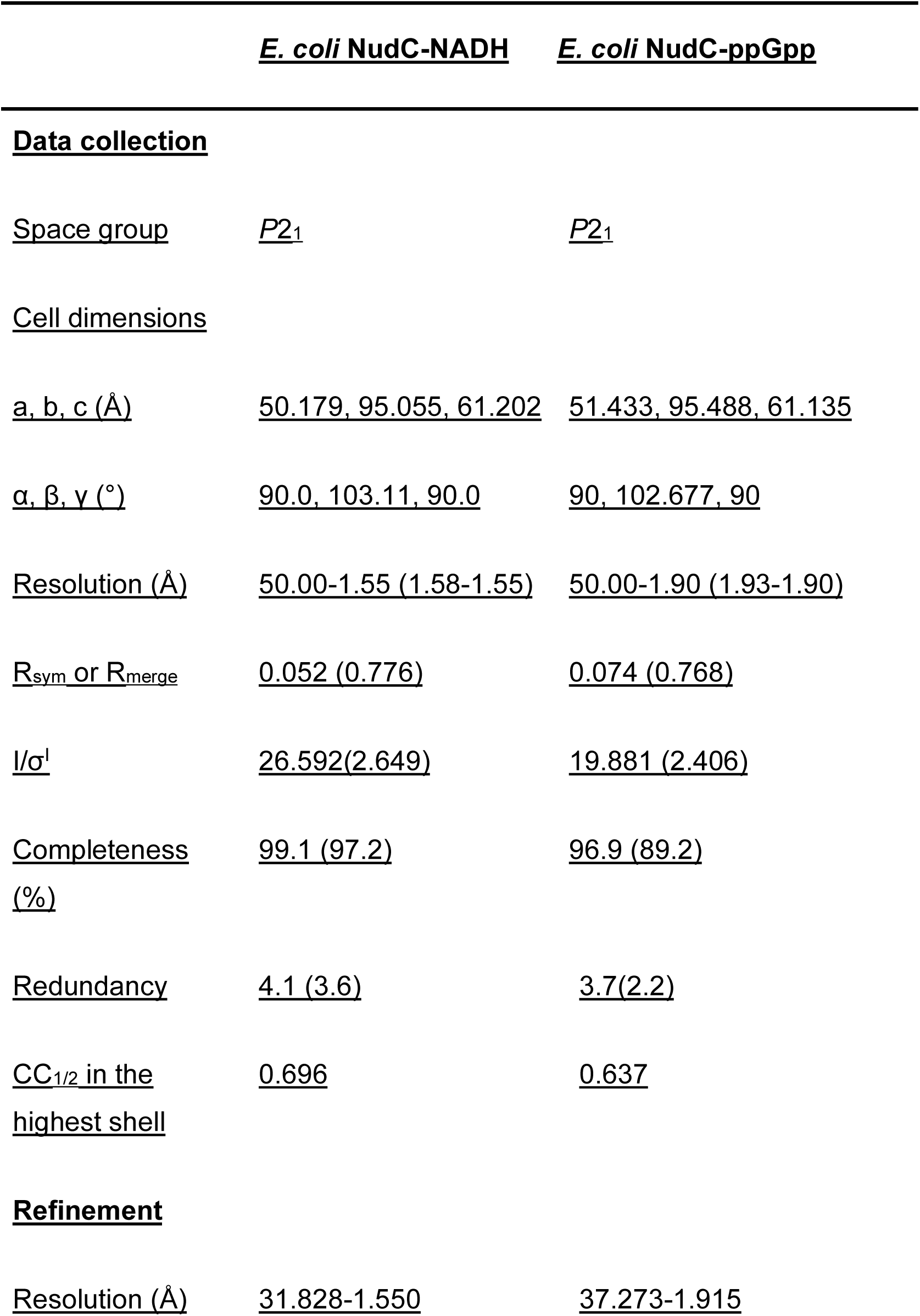
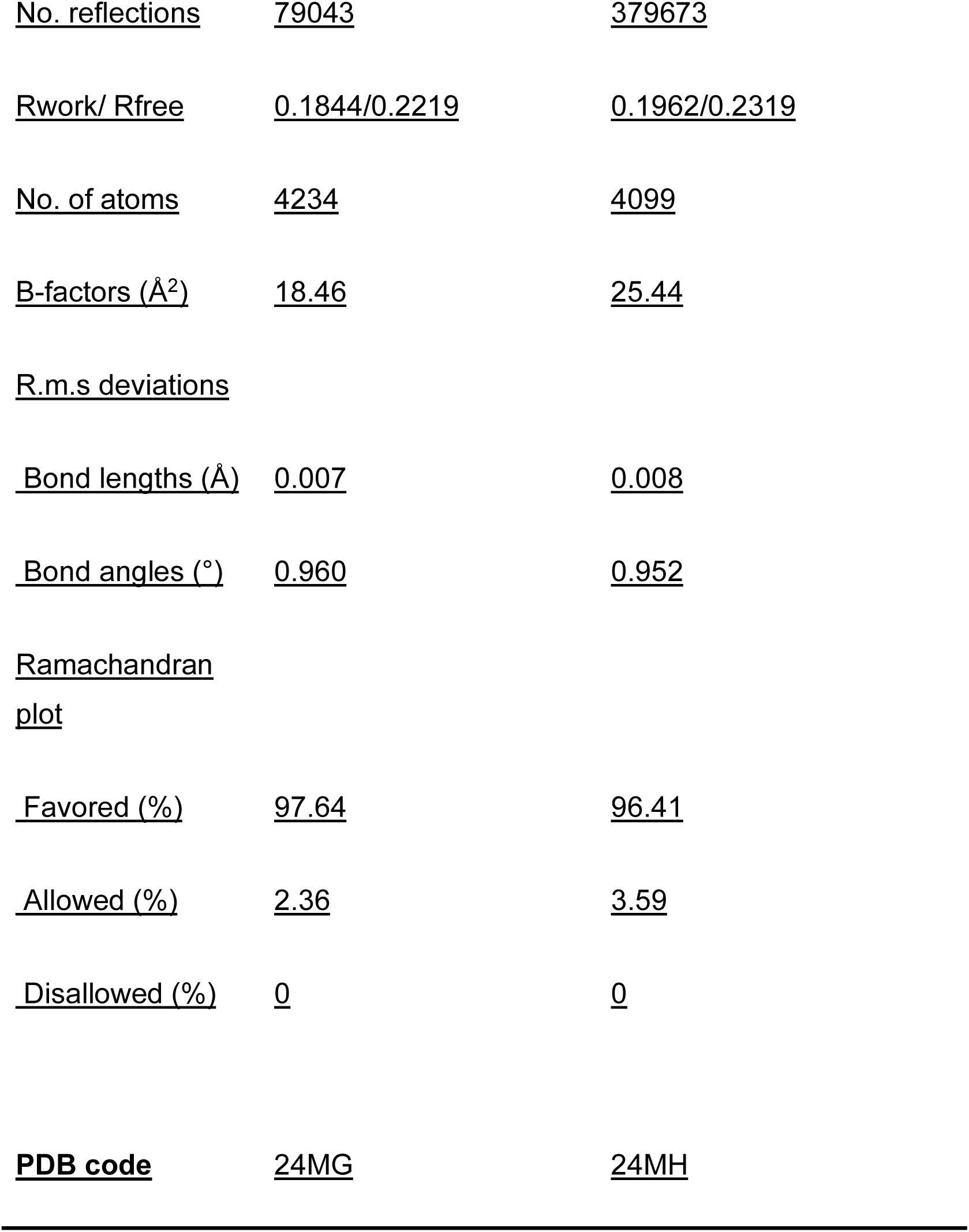
Crystal statistics of *E. coli* NudC-NADH and *E. coli* NudC-ppGpp.

## MATERIALS AND REAGENTS

**Supplementary Table 2:**
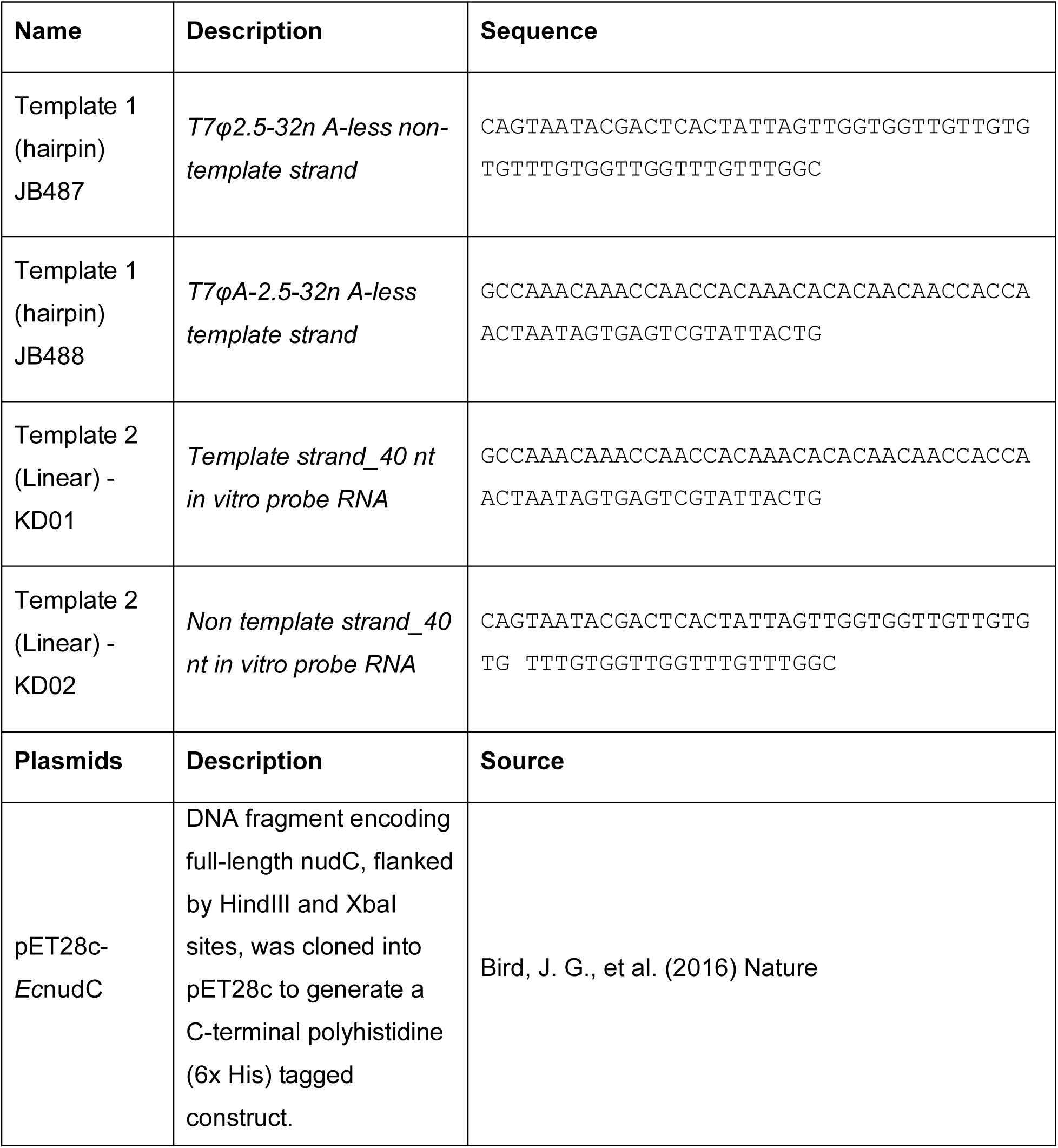
Oligonucleotides used in this study.

**Supplementary Table 3:** Reagents used in this study.

| **Reagent** | **Source** | **Catalog number** |
|---|---|---|
| [α32P]-GTP | Revvity | BLU506H250UC |
| Ammonium persulfate | Fisher Scientific | BP179-100 |
| Ultrapure rNTP set | Cytiva | GE27-2025-01 |
| Acid Phenol/Chloroform/Isoamyl alcohol 125:24:1 | Sigma Aldrich | P3803-400ML |
| PBA (3′-acrylamidophenylboronic acid) | Boron Molecular, Inc. | BM1195 |
| Phusion HF Master Mix | Fisher Scientific | F531S |
| QIAquick PCR purification/cleanup kit | Qiagen | 28106 |
| SequaGel UreaGel sequencing system | National Diagnostics | EC-833 |
| 3M Sodium acetate, trihydrate (NaOAc) | Fisher Scientific | AM9740 |
| T7 RNA Polymerase | New England Biolabs | M0251S |
| Ethanol (200 Proof) | Decon Labs | 2701 |
| Ambion Nuclease-free water | Fisher Scientific | AM9932 |
| TAE (Tris acetate EDTA) solution | Genesee Scientific | 18-134L |
| TEMED | Fisher Scientific | 50-197-8483 |
| Tris base | Fisher Scientific | BP1525 |
| NAD<sup>+</sup> (grade I, free acid) | Sigma-Aldrich | 10127965001 |
| NADH (grade I, free acid) | Sigma-Aldrich | 10107735001 |
| ATP | GoldBio | A-081-1 |
| Isopropyl β-D-1- thiogalactopyranoside (IPTG) | GoldBio | I2481C25 |
| Ni-NTA Agarose | MCLAB | NINTA-25 |
| Amicon Ultra Centrifugal Filter, 10kDa MWCO | Sigma Aldrich | UFC903024 |
| NuPAGE LDS sample buffer | Thermo Fisher Scientific | NP0007 |
| Pall Nanosep columns, 10k cutoff 24/pk | Fisher Scientific | 50-197-9585 |
| Illustra MicroSpin G-25 columns | Cytiva | 27532501 |
| Whatman Vacu-guard filter | Fisher Scientific | 09-744-75 |
| NOVEX 4-12% Bis-Tris polyacrylamide gels | ThermoFisher |  |
| Saran/polyethylene wrap | SC Johnson |  |

**Supplementary Table 4:**
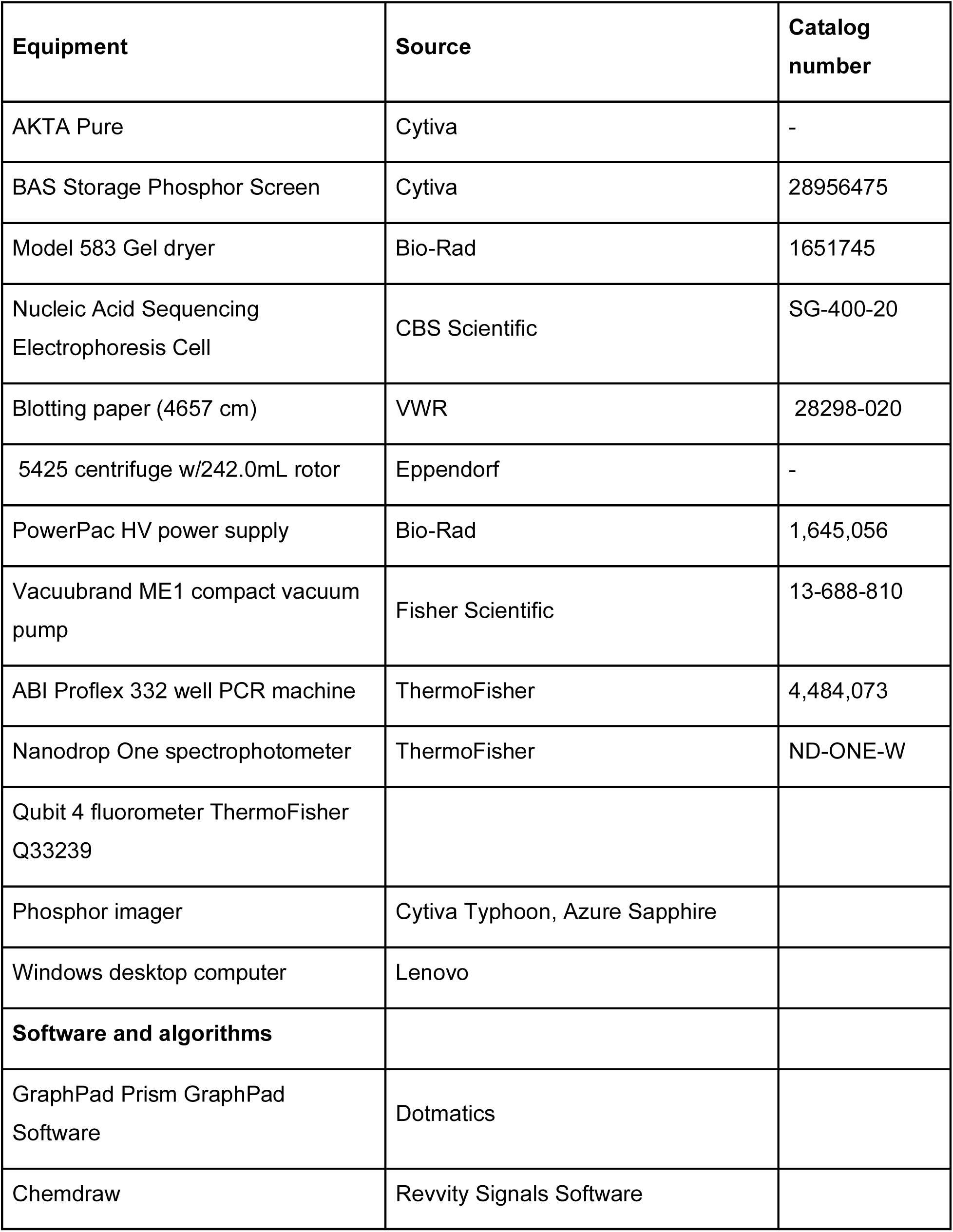
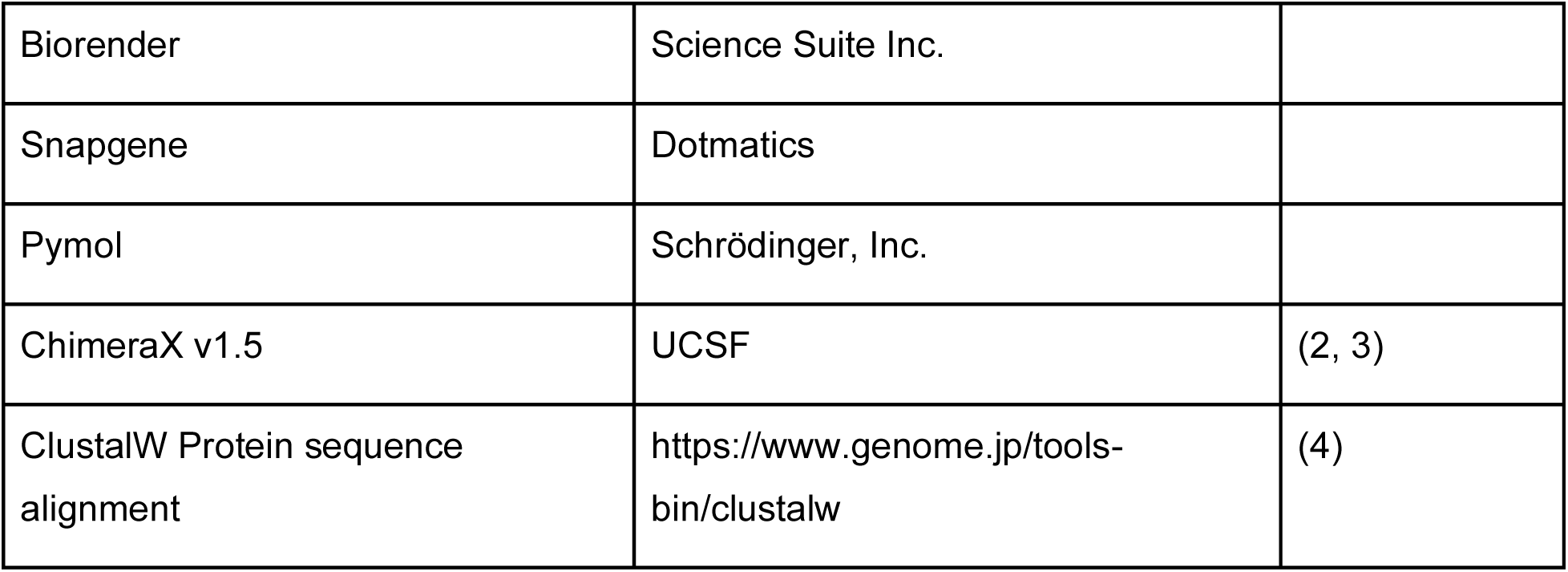
Equipment and software used in this study.

**Supplementary Table 5:** Buffer compositions.

| **Buffers** | **Composition** |
|---|---|
| RNA Polymerase buffer | 50mM Tris, pH 8.0<br>2mM Spermidine<br>5mM MgCl2<br>1mM DTT |
| Buffer 2 | 10mM Tris, pH 8.0<br>50mM NaCl<br>10mM MgCl2<br>1mM DTT |
| STOP solution | 50mM Tris, pH 8.0<br>18mM EDTA<br>0.1mg/mL Glycogen |
| Transcription loading buffer | 0.5 TBE<br>90% Formamide<br>18mM EDTA<br>0.2% SDS<br>0.05% Bromophenol Blue<br>0.05% Xylene Cyanol<br>0.05% Amaranth Red (optional) |

## Notes

### Competing Interest Statement

The authors have declared no competing interest.

## References

1. Chen, Y.G., Kowtoniuk, W.E., Agarwal, I., et al. (2009) LC/MS analysis of cellular RNA reveals NAD-linked RNA. Nat. Chem. Biol., 5, 879–881.

2. Kowtoniuk, W.E., Shen, Y., Heemstra, J.M., et al. (2009) A chemical screen for biological small molecule–RNA conjugates reveals CoA-linked RNA. Proc. Natl. Acad. Sci., 106, 7768–7773.

3. Wang, J., Alvin Chew, B.L., Lai, Y., et al. (2019) Quantifying the RNA cap epitranscriptome reveals novel caps in cellular and viral RNA. Nucleic Acids Res., 47, e130.

4. Cahová, H., Winz, M.-L., Höfer, K., et al. (2015) NAD captureSeq indicates NAD as a bacterial cap for a subset of regulatory RNAs. Nature, 519, 374–377.

5. Walters, R.W., Matheny, T., Mizoue, L.S., et al. (2017) Identification of NAD+ capped mRNAs in Saccharomyces cerevisiae. Proc. Natl. Acad. Sci., 114, 480–485.

6. Jiao, X., Doamekpor, S.K., Bird, J.G., et al. (2017) 5’ End Nicotinamide Adenine Dinucleotide Cap in Human Cells Promotes RNA Decay through DXO-Mediated deNADding. Cell, 168, 1015–1027.e10.

7. Ruiz-Larrabeiti, O., Benoni, R., Zemlianski, V., et al. (2021) NAD+ capping of RNA in Archaea and Mycobacteria. BioRxiv.

8. Gomes-Filho, J.V., Breuer, R., Morales-Filloy, H.G., et al. (2023) Identification of NAD-RNA species and ADPR-RNA decapping in Archaea. Nat. Commun., 14, 7597.

9. Wolfram-Schauerte, M., Keuthen, H., Moskalchuk, A., et al. (2026) T4 phage RNA is NAD-capped and alters the NAD-cap epitranscriptome of Escherichia coli during infection. 10.1101/2024.04.04.588121.

10. Sherwood, A.V., Rivera-Rangel, L.R., Ryberg, L.A., et al. (2023) Hepatitis C virus RNA is 5’-capped with flavin adenine dinucleotide. Nature, 619, 811–818.

11. Bird, J.G., Zhang, Y., Tian, Y., et al. (2016) The mechanism of RNA 5’ capping with NAD +, NADH and desphospho-CoA. Nature, 535, 444–447.

12. Zhang, H., Zhong, H., Wang, X., et al. (2021) Use of NAD tagSeq II to identify growth phase-dependent alterations in E. coli RNA NAD+ capping. Proc. Natl. Acad. Sci., 118, e2026183118.

13. Mackie, G.A. (1998) Ribonuclease E is a 5’-end-dependent endonuclease. Nature, 395, 720–724.

14. Celesnik, H., Deana, A. and Belasco, J.G. (2007) Initiation of RNA Decay in Escherichia coli by 5’ Pyrophosphate Removal. Mol. Cell, 27, 79–90.

15. Deana, A., Celesnik, H. and Belasco, J.G. (2008) The bacterial enzyme RppH triggers messenger RNA degradation by 5’ pyrophosphate removal. Nature, 451, 355–358.

16. Callaghan, A.J., Marcaida, M.J., Stead, J.A., et al. (2005) Structure of Escherichia coli RNase E catalytic domain and implications for RNA turnover. Nature, 437, 1187–1191.

17. Gao, A., Vasilyev, N., Kaushik, A., et al. (2020) Principles of RNA and nucleotide discrimination by the RNA processing enzyme RppH. Nucleic Acids Res., 48, 3776–3788.

18. Höfer, K., Li, S., Abele, F., et al. (2016) Structure and function of the bacterial decapping enzyme NudC. Nat. Chem. Biol., 12, 730–734.

19. Zhou, W., Guan, Z., Zhao, F., et al. (2021) Structural insights into dpCoA-RNA decapping by NudC. RNA Biol., 18, 244–253.

20. Zhang, D., Liu, Y., Wang, Q., et al. (2016) Structural basis of prokaryotic NAD-RNA decapping by NudC. Cell Res., 26, 1062–1066.

21. Frick, D.N. and Bessman, M.J. (1995) Cloning, Purification, and Properties of a Novel NADH Pyrophosphatase: EVIDENCE FOR A NUCLEOTIDE PYROPHOSPHATASE CATALYTIC DOMAIN IN MutT-LIKE ENZYMES (*). J. Biol. Chem., 270, 1529–1534.

22. Bouvier, M. and Carpousis, A.J. (2011) A tale of two mRNA degradation pathways mediated by RNase E. Mol. Microbiol., 82, 1305–1310.

23. Frindert, J., Zhang, Y., Nübel, G., et al. (2018) Identification, Biosynthesis, and Decapping of NAD-Capped RNAs in B. subtilis. Cell Rep., 24, 1890–1901.e8.

24. Wang, B., Dai, P., Ding, D., et al. (2019) Affinity-based capture and identification of protein effectors of the growth regulator ppGpp. Nat. Chem. Biol., 15, 141–150.

25. Gatewood, M.L. and Jones, G.H. (2010) (p)ppGpp Inhibits Polynucleotide Phosphorylase from Streptomyces but Not from Escherichia coli and Increases the Stability of Bulk mRNA in Streptomyces coelicolor. J. Bacteriol., 192, 4275–4280.

26. Hsieh, P., Richards, J., Liu, Q., et al. (2013) Specificity of RppH-dependent RNA degradation in Bacillus subtilis. Proc. Natl. Acad. Sci., 110, 8864–8869.

27. Bennett, B.D., Kimball, E.H., Gao, M., et al. (2009) Absolute metabolite concentrations and implied enzyme active site occupancy in Escherichia coli. Nat. Chem. Biol., 5, 593–599.

28. Hauryliuk, V., Atkinson, G.C., Murakami, K.S., et al. (2015) Recent functional insights into the role of (p)ppGpp in bacterial physiology. Nat. Rev. Microbiol., 13, 298–309.

29. Potrykus, K. and Cashel, M. (2008) (p)ppGpp: Still Magical?*. Annu. Rev. Microbiol., 62, 35–51.

30. Vvedenskaya, I.O., Bird, J.G., Zhang, Y., et al. (2018) CapZyme-Seq Comprehensively Defines Promoter-Sequence Determinants for RNA 5’ Capping with NAD+. Mol. Cell, 70, 553–564.e9.

31. Bird, J.G., Basu, U., Kuster, D., et al. (2018) Highly efficient 5’ capping of mitochondrial RNA with NAD+ and NADH by yeast and human mitochondrial RNA polymerase. eLife, 7, e42179.

32. Mickutė, M., Krasauskas, R., Kvederavičiūtė, K., et al. (2023) Interplay between bacterial 5’-NAD-RNA decapping hydrolase NudC and DEAD-box RNA helicase CsdA in stress responses. mSystems, 8, e00718-23.

33. Luciano, D.J. and Belasco, J.G. (2015) NAD in RNA: unconventional headgear. Trends Biochem. Sci., 40, 245–247.

34. Liu, W., Hu, K., Nie, S., et al. (2026) NAD+ capping of sibD transcripts in E. coli is mediated by its minimal promoter and enhanced by ppGpp. Nucleic Acids Res., 54, gkag102.

35. Singh, Y. and Bird, J.G. (2022) Chapter Thirteen - A gel electrophoresis-based assay for measuring enzymatic RNA decapping activity. In Shukla, A.K. (ed), Integrated Methods in Protein Biochemistry: Part A, Methods in Enzymology. Academic Press, Vol. 675, pp. 323–350.

36. Nübel, G., Sorgenfrei, F.A. and Jäschke, A. (2017) Boronate affinity electrophoresis for the purification and analysis of cofactor-modified RNAs. Methods, 117, 14–20.

## References

1. Zhang, Y., Gong, C., Sun, J., et al. (2026) Protenix-v2: Broadening the Reach of Structure Prediction and Biomolecular Design. 10.64898/2026.04.10.717613.

2. Meng, E.C., Goddard, T.D., Pettersen, E.F., et al. (2023) UCSF ChimeraX: Tools for structure building and analysis. Protein Science, 32, e4792.

3. Pettersen, E.F., Goddard, T.D., Huang, C.C., et al. (2021) UCSF ChimeraX: Structure visualization for researchers, educators, and developers. Protein Science, 30, 70–82.

4. Thompson, J.D., Higgins, D.G. and Gibson, T.J. (1994) CLUSTAL W: improving the sensitivity of progressive multiple sequence alignment through sequence weighting, position-specific gap penalties and weight matrix choice. Nucleic Acids Res, 22, 4673–4680.

